# GeroQubit: a lightweight, honesty-first de-novo design platform for geroscience-native small molecules with calibrated uncertainty

**DOI:** 10.64898/2026.06.07.730687

**Authors:** Dinesh K, H. Swetha

## Abstract

Computational molecule generation has outpaced its own credibility. We present GeroQubit, a GPU-free de-novo design platform that organizes candidates along a target × tissue × hallmark model and reports every signal alongside its measured baseline. We treat our tissue aging-signature readout as a mechanistic structural prior that we explicitly disclose is *not* validated against lifespan, and we surface efficacy only through a structure-to-lifespan k-NN whose weak but real signal (leave-one-out ρ ≈ 0.145) is wrapped in empirically-calibrated conformal intervals (90% target, 90.3% measured coverage). On a held-out retrospective recovery of ∼1,940 ChEMBL binders against decoys, the score reaches ROC-AUC 0.945 with ∼20× enrichment at 1% (BEDROC 0.91) and survives a scaffold-disjoint split — yet we report that it collapses to near-random (AUC 0.62) on genuinely novel chemotypes. Molecules are assembled reaction-first, so every candidate carries a verified synthetic route and atom-level synthon provenance; ADMET is handled as a multi-objective Pareto problem. We frame the disclosed weak signals and the hard-case failures not as flaws but as the honest, decision-useful output the field’s own critics demand.

## 1. Introduction

Computational molecule generation has outpaced its own credibility: aging clocks are correlative, uncertainty is rarely reported, and benchmark scores seldom predict potency. GeroQubit occupies the thin white space of lightweight, GPU-free, small-molecule geroscience design, under a measure-or-don’t-ship discipline. We argue that honest uncertainty is a competitive feature, not an apology.

## 2. The design model — target × tissue × hallmark

Candidates are reasoned about across validated longevity targets, multiple tissues, and the twelve hallmarks of aging simultaneously, because aging is multi-causal and single-tissue framing is misleading. This factored representation lets one molecule be evaluated for systemic breadth and rational combination, rather than against a single endpoint.

## 3. Phenotype-level scoring (mechanistic prior)

Each structure is projected into a 128-dimensional transcriptomic embedding and scored for its predicted perturbation of a curated eight-gene aging signature (NAD^+^ axis, senescence, inflammaging) across ten tissues. We present this strictly as a mechanistic structural prior for prioritization; it is not lifespan-validated and never enters a selection objective as an efficacy claim.

## 4. Efficacy priors & honest uncertainty

The only efficacy signal we report is a structure-to-measured-lifespan k-nearest-neighbour regression over a curated longevity compound set, accompanied by its disclosed leave-one-out correlation. Each prediction ships with a normalized inductive conformal interval that widens for out-of-domain or disagreeing-neighbour molecules, with empirically verified coverage. Wide intervals are shown, not hidden.

## 5. De-novo assembly & synthesizability

Molecules are built through verified reaction steps rather than emitted as raw strings, so every candidate has a synthesizable route by construction. We attach atom-level synthon provenance — true synthesis attribution rather than post-hoc explainability theatre.

## 6. Multi-objective ADMET (Pareto)

Drug-likeness, ADMET endpoints, and complexity are treated as competing objectives on a Pareto frontier relevant to chronic dosing, rather than collapsed into one opaque score. Applied at final selection (not folded into search), this preserves the potency champion while surfacing the ADMET-clean tail — peak fitness identical across 10/10 targets in a same-pool test, with ADMET desirability up +0.02–0.08.

## 7. RESULTS

### Three candidates against mTORC1 — complete output, real values

From 416 generated molecules across whole-body mTORC1 runs, ADMET-filtered, we show three representative candidates spanning the drug-likeness / activity trade-off. Every number below is the platform’s actual output, including the disclosed limitations.

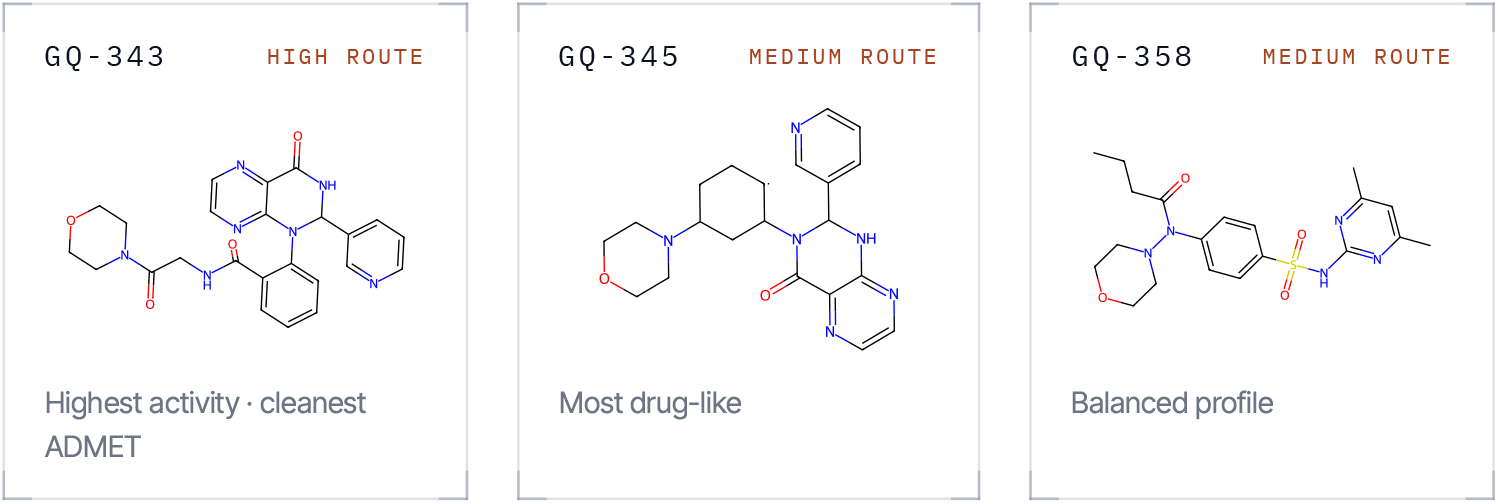

## 8. VALIDATION

**TABLE 1.**
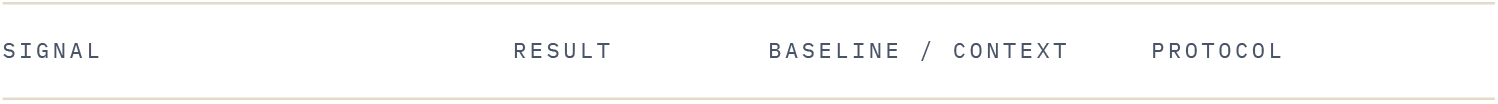
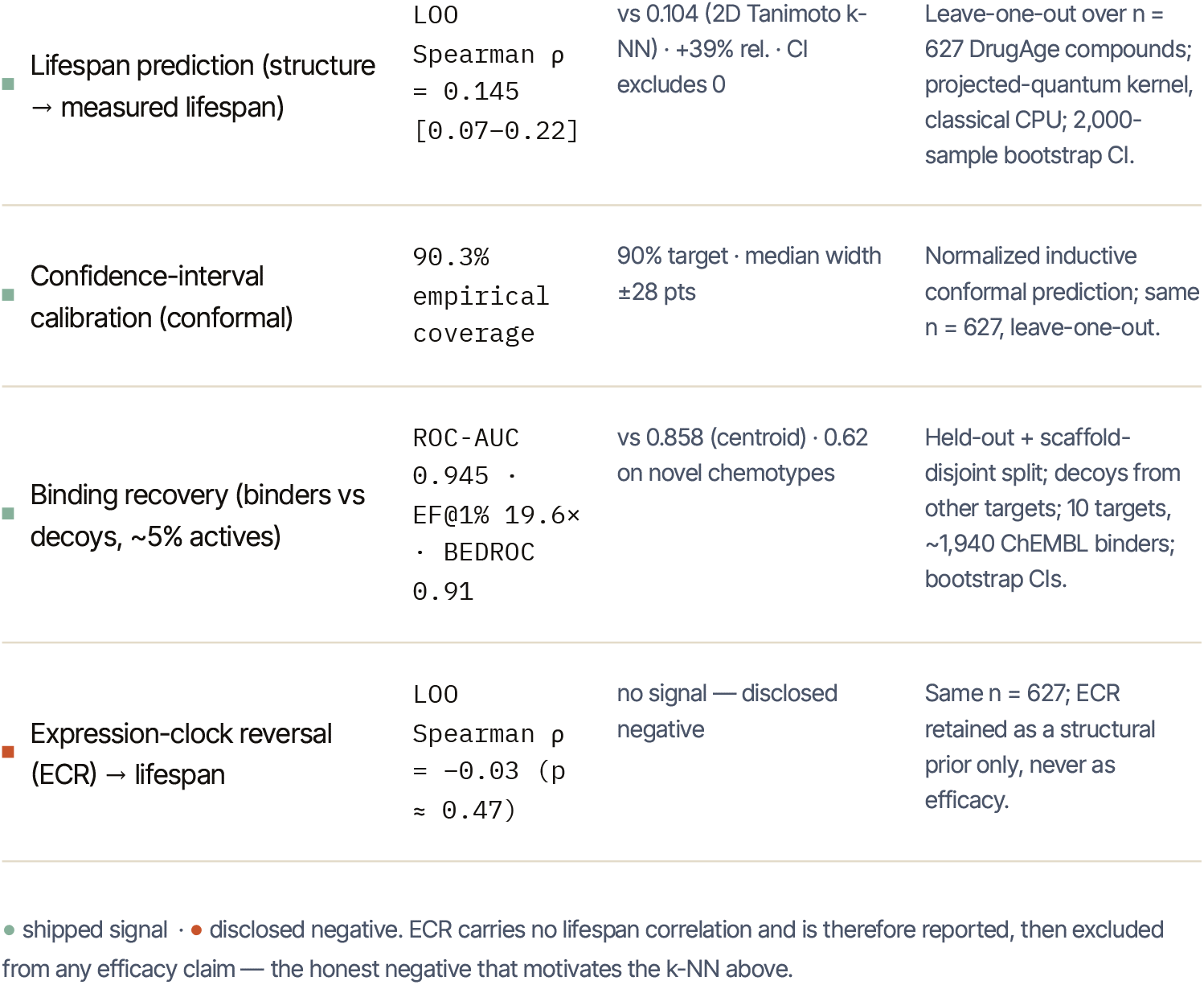
EVERY SIGNAL VS ITS BASELINE, ON HELD - OUT DATA.

**TABLE 2.**
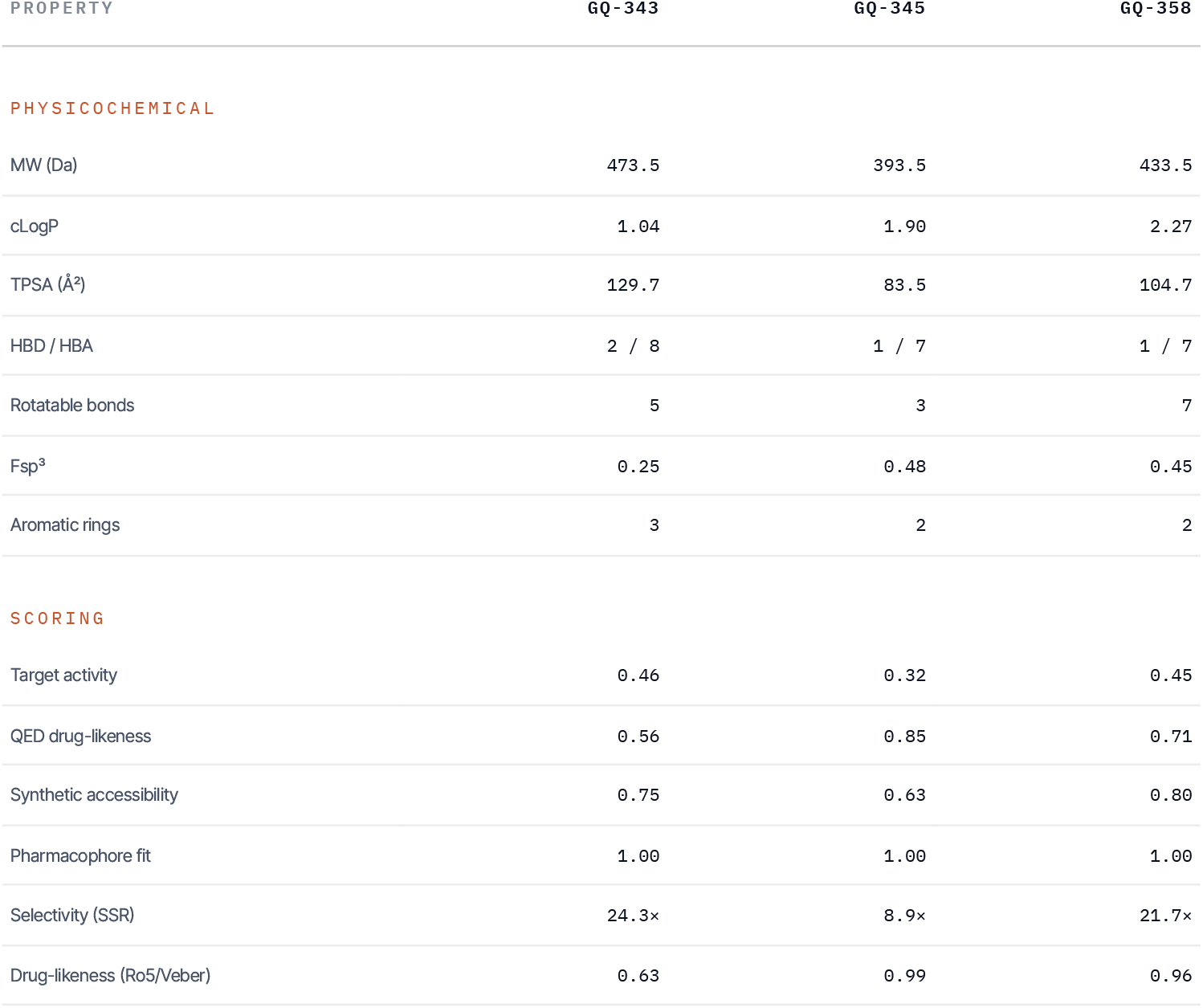
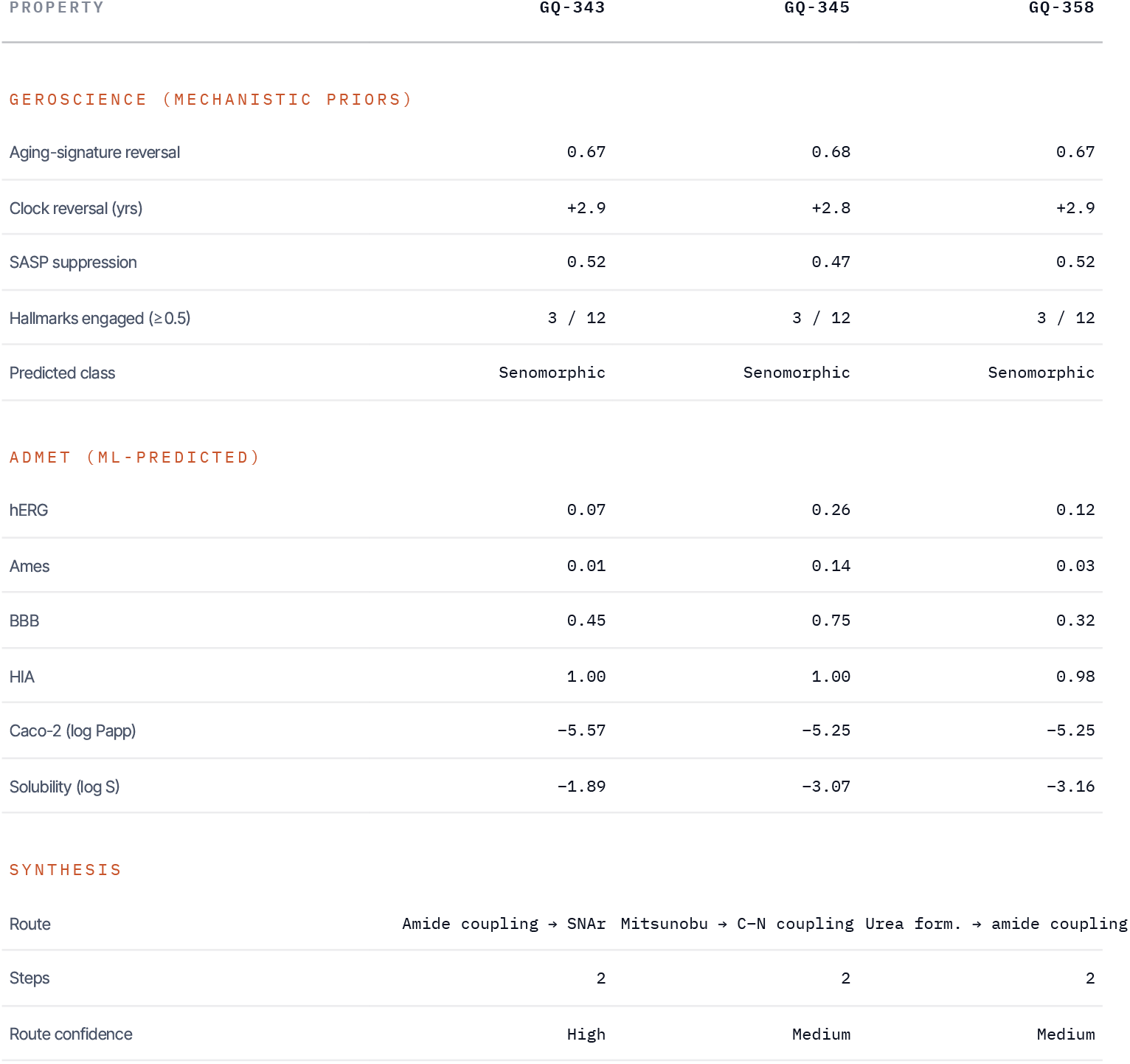
THREE REPRESENTATIVE CANDIDATES, COMPLETE PLATFORM OUTPUT (REAL VALUES)

> “We publish the disclosed negative because it is the control that makes the positive credible: a weak-but-real lifespan signal, honestly bounded, is worth more than a strong number nobody can reproduce.”

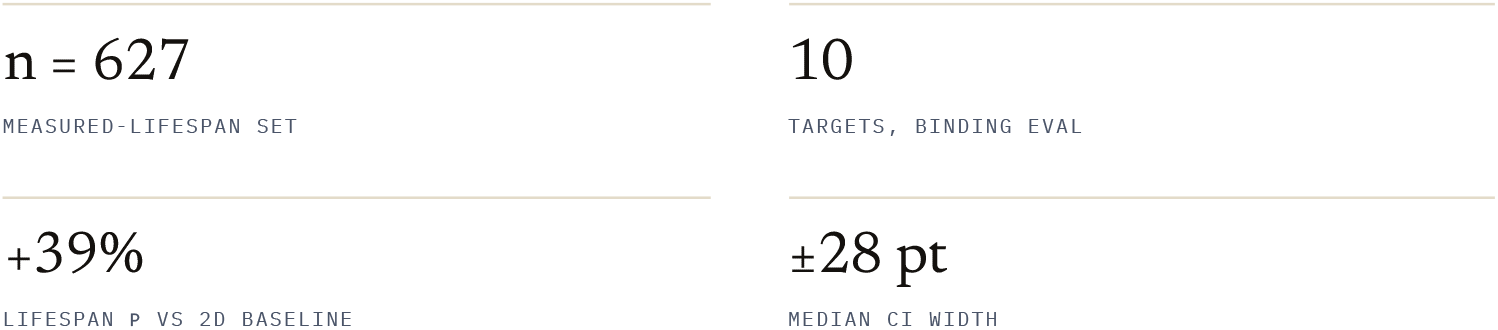

### 8B · RETROSPECTIVE RECOVERY & BENCHMARK

#### DOES IT PULL KNOWN ACTIVES BACK OUT?

Beyond a single AUC, we ran a standard virtual-screening recovery: hide each target’s known binders in a realistic decoy pool (∼5% actives) and measure early-recognition enrichment, with bootstrap confidence intervals — and report the hard cases (unseen scaffolds, novel chemotypes) next to the easy ones.

**TABLE 2.** HELD - OUT RECOVERY OF KNOWN BINDERS VS DECOYS (10 TARGETS, ∼5% ACTIVE FRACTION)

| REGIME | ROC-AUC [95% CI] | EF@1% | BEDROC |
| --- | --- | --- | --- |
| ■ Random split (standard) | 0.945 [0.93–0.96] | 19.6× | 0.91<br>[0.89–0.93] |
| ■ Scaffold-disjoint split | 0.945 [0.93–0.96] | 21.1× | 0.90<br>[0.88–0.92] |
| ■ Novel chemotype (max-Tan < 0.35) | 0.62 | ≈ 0 | — |
| ■ Centroid baseline (random split) | 0.858 | — | 0.75 |

● strong recovery · ● honest hard case. Enrichment is real (EF@1% ≈ 20× over random) and survives a scaffold-disjoint split — but **collapses to near-random on genuinely 2D-novel chemotypes** (AUC 0.62). We report that ceiling: a 2D similarity score cannot generalise to scaffolds unlike anything it has seen, and we don't pretend it can.

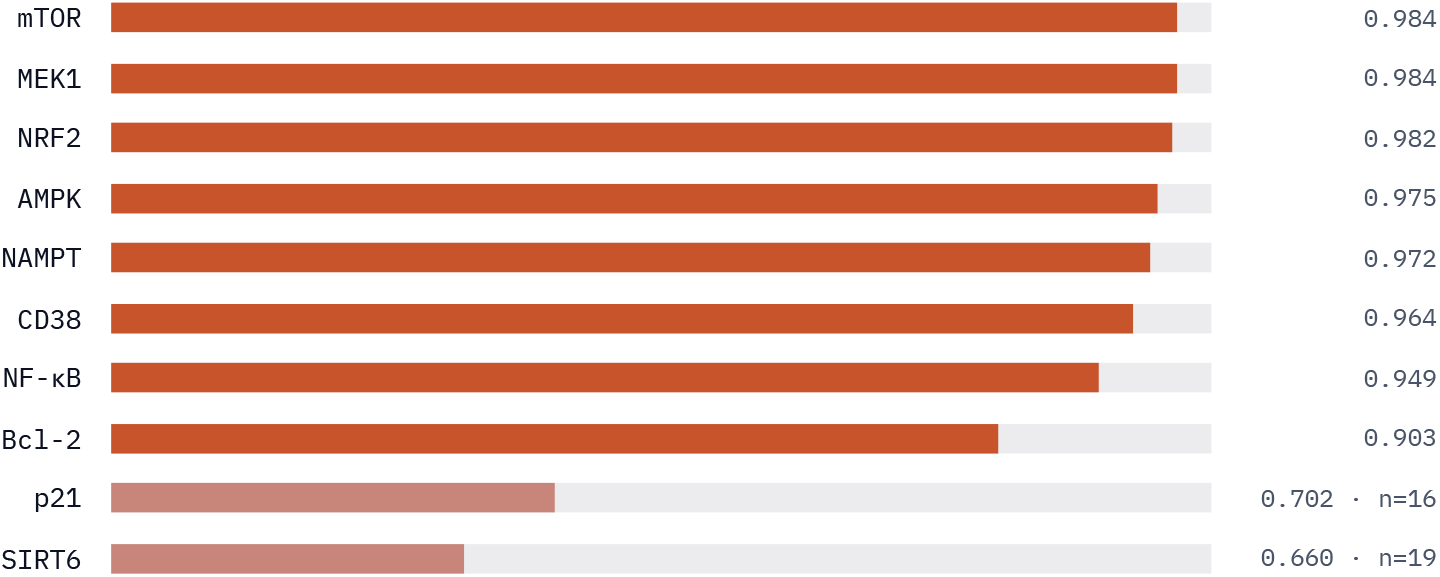

Eight of ten targets recover strongly (AUC 0.90–0.98; mTOR 0.98, Bcl-2 0.90). The two weak targets are the two with the fewest known binders (SIRT6 n=19, p21 n=16) — a **data-scarcity** limit, disclosed, not a modelling claim. Bars scaled from 0.5 (random).

**TABLE 3.** PLATFORM METHOD VS THE HONEST BASELINE IT MUST BEAT.

|  |  |  |  |
| --- | --- | --- | --- |
| Lifespan rank (LOO Spearman ρ) | Projected-quantum-kernel k-NN — 0.145<br>[0.07–0.22] | Tanimoto k-NN — 0.104<br>[0.03–0.18] | +39% |
| Binding recovery (pooled ROC-AUC) | k-NN-max to binders — 0.945 | Tanimoto-to-centroid — 0.858 | +0.087 |
| Binding early recognition (BEDROC) | k-NN-max — 0.91 | centroid — 0.75 | +0.16 |

The discipline: a method ships only if it beats the strongest honest baseline. The quantum kernel's +39% on lifespan rank is real (bootstrap CI excludes zero) but the signal is weak in absolute terms — so it ships *with* its conformal intervals, never as a promise.

### 8C · GENETICS – ANCHORED TARGET PRIORITISATION

#### WHICH TARGETS DOES HUMAN GENETICS ACTUALLY BACK?

The strongest single predictor of clinical success is human genetic causal evidence (∼2× approval odds; Nelson 2015). We score each target by a **Human Causal Aging Evidence (HCAE)** index — the Open Targets human genetic association to age-related diseases, weighted by small-molecule druggability — and rank the platform’s targets by it. The honest result: **four of the ten original targets (SIRT6, NAMPT, AMPK, NRF2) carry zero human genetic aging evidence** — they rest on model-organism and mechanistic evidence only. We disclose this rather than bury it.

| TARGET | HCAE | AGE-RELATED DISEASES | DRUGGABLE |
| --- | --- | --- | --- |
| ■ p21 (CDKN1A) | 0.31 | 5 | 0.3 |
| ■ IGF1R | 0.19 | 2 | 1.0 |
| ■ CD38 | 0.13 | 2 | 0.55 |
| ■ Bcl-2 | 0.11 | 1 | 1.0 |
| ■ mTOR | 0.08 | 1 | 1.0 |
| ■ MEK1 | 0.02 | 1 | 1.0 |
| ■ NF-κB | 0.01 | 0 | 0.9 |
| ■ SIRT6 | 0.00 | 0 | 0.55 |
| ■ NAMPT | 0.00 | 0 | 0.9 |
| ■ AMPK | 0.00 | 0 | 0.9 |
| ■ NRF2 | 0.00 | 0 | 1.0 |

● human-genetics-backed · ● model-organism evidence only. This ranking now drives target selection in the platform. It also surfaced the right expansion: we added **IGF1R** — the canonical IGF-1/insulin (daf-2) longevity axis, genetics-backed and fully druggable (276 ChEMBL binders) — as the 11th target. The top senescence-genetics gene, CDKN2A/p16, was *excluded*: it has zero small-molecule druggability and no ChEMBL binders, which the druggability gate caught automatically.

Disclosed future work: a drug-target Mendelian-randomisation layer (cis-eQTL/pQTL instruments → human lifespan/healthspan outcomes) would upgrade this from associative to causal-and-directional, with most targets expected to return an honest null.

**FIG. 1.**
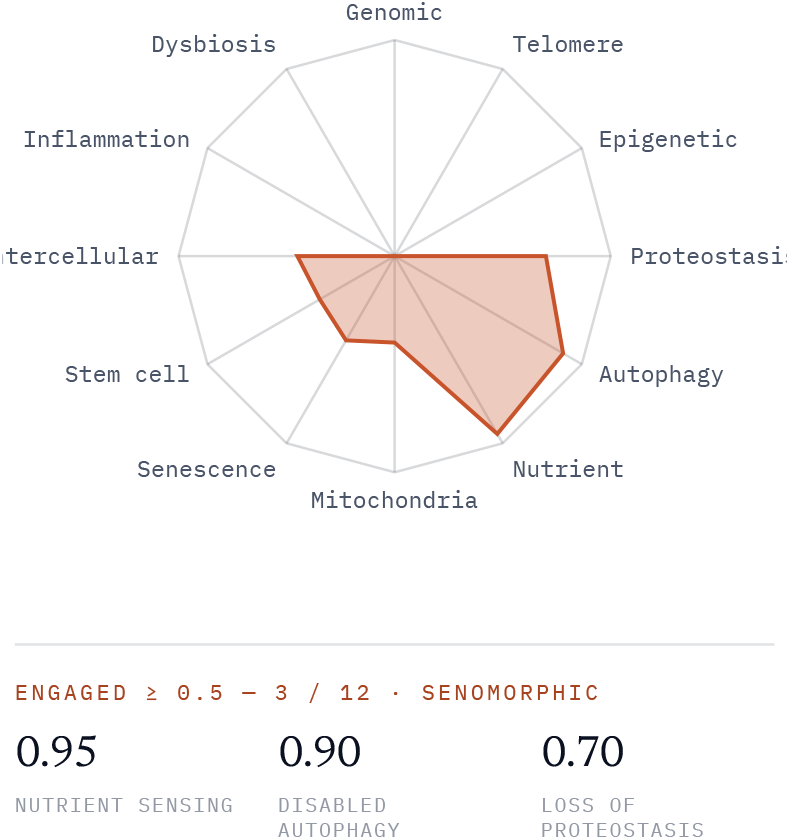
HALLMARK ENGAGEMENT (LÓPEZ-OTÍN12), CANDIDATEGQ - 259

### PER - CANDIDATE, NOT PER - HEADLINE

#### Every molecule reports its own hallmark profile

Breadth is treated as biology, not a single number. The same radar shown here is produced for every candidate inside the platform and exported with the dossier — so a reviewer sees exactly which aging programs a molecule is predicted to engage, and how strongly.

GQ-259 concentrates on the nutrient-sensing / autophagy / proteostasis axis — the classic mTOR-linked senomorphic signature — while leaving the genomic, telomere, and epigenetic programs untouched. That selectivity is the point: an honest profile, not a uniform “engages everything” claim.

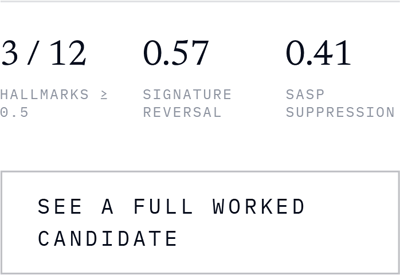

### GENE - LEVEL PHENOTYPE

#### A 128-D embedding, an 8-gene signature

Each structure is projected into a 128-dimensional transcriptomic embedding and scored for its predicted Δ on a curated eight-gene aging panel — the NAD+ axis (NAMPT, CD38, SIRT6), senescence (CDKN2A, BCL2), inflammaging (IL6, TNF), and mTOR. A mechanistic prior, disclosed — not validated against expression or lifespan.

**FIG. 2.**
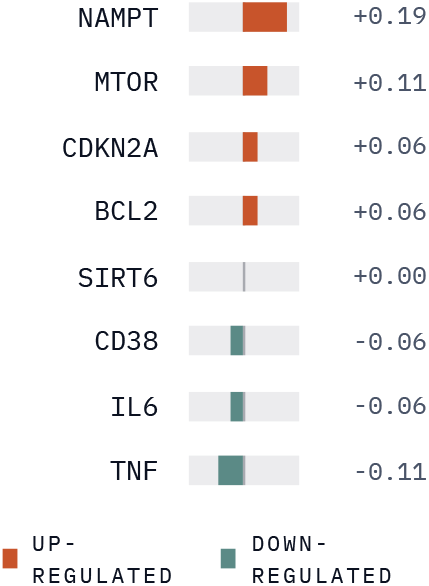
8 - GENE AGING SIGNATURE · GQ - 259

## 9 · LIMITATIONS

The tissue aging-signature / clock-reversal readout is a mechanistic structural prior only and is **not** validated against lifespan (leave-one-out ρ ≈ −0.03); it is used for prioritization, never as an efficacy claim. The single efficacy signal we report, a structure-to-lifespan k-NN, is genuinely **weak** (ρ ≈ 0.145), and we therefore publish its conformal intervals, which are correspondingly wide. Binding-plausibility AUC is **retrospective**, measured on known binders versus decoys, not prospective affinity — which honestly requires GPU/physics outside our lightweight scope. Where 3D is used it relies on a single conformer. All outputs are computational hypotheses requiring experimental confirmation; the platform is research-use only and not for clinical, diagnostic, or therapeutic decisions.

### HONEST ANSWERS

#### The questions a careful buyer asks first

##### Is your aging-clock / phenotype reversal validated against lifespan?

No — and we say so up front. We audited our expression-clock-reversal (ECR) signal against 627 DrugAge compounds with measured lifespan extension and found no correlation (leave-one-out Spearman ρ ≈ −0.03, not significant). Clock-reversal and the hallmark/tissue phenotype are therefore used only as mechanistic structural priors for prioritisation — never as an efficacy claim. The one efficacy signal we report is a structure→measured-lifespan k-NN (LOO ρ ≈ 0.145, +39% over a 2D baseline, bootstrap CI excludes zero), shipped with a calibrated conformal interval. A weak but honest prior, disclosed with its limits.

##### Isn’t a 0.945 recovery AUC just an easy retrospective artifact?

Partly — which is exactly why we stress-test it. Recovery stays strong under a scaffold-disjoint split (AUC 0.945, EF@1% ≈ 20×, BEDROC 0.91), but we also publish the regime where it fails: on genuinely 2D-novel chemotypes (max-Tanimoto < 0.35 to the reference) it collapses to near-random (AUC 0.62). The score is a strong 2D recovery engine and a poor de-novo extrapolator, and we report both numbers rather than only the flattering one.

##### Does “quantum” mean you need a quantum computer?

No. The lifespan predictor uses a projected quantum kernel (Huang et al., Nat. Commun. 2021) — a classically computable similarity that is non-linear in substructure overlap and runs in a single matrix multiply on an ordinary CPU. No GPU, no quantum hardware. It is the one place the quantum-inspired idea earns its keep: a measured +39% over plain Tanimoto on our single validated signal.

## DECLARATIONS

### AUTHOR CONTRIBUTIONS

Dinesh K and H Swetha jointly conceived the platform, designed the scoring and validation methodology, performed the analyses, and wrote the manuscript. Both authors reviewed and approved the final version.

### DATA & CODE AVAILABILITY

Validation draws on public datasets — DrugAge (lifespan), ChEMBL (bioactivity), GTEx (tissue expression), and the López-Otín hallmark framework. Aggregate validation results are reported in full in this manuscript. The platform’s internals — scoring weights, the transcriptomic embedding, generator encoding, and reaction templates — are proprietary and not disclosed. Representative candidate structures are available from the corresponding author under a research agreement.

### ETHICS

No human or animal subjects were involved; all results are computational. The platform is a research tool for hypothesis generation and is not intended for clinical, diagnostic, or therapeutic use.

### PREPRINT STATUS

This manuscript is a preprint and has not been peer-reviewed. A reuse license is selected at the time of posting.

